# Radiator: a cloud-based framework for deploying re-usable bioinformatics tools

**DOI:** 10.1101/614594

**Authors:** Emily K.W. Lo, Remy M. Schwab, Zak Burke, Patrick Cahan

**Affiliations:** Institute for Cell Engineering, Johns Hopkins University, Baltimore, MD, 21205, USA; Department of Biomedical Engineering, Johns Hopkins University, Baltimore, MD, 21205, USA

## Abstract

**Summary:** Accessibility and usability of compute-intensive bioinformatics tools can be increased with simplified web-based graphic user interfaces. However, deploying such tools as web applications presents additional barriers, including the complexity of developing a usable interface, network latency in transferring large datasets, and cost, which we encountered in developing a web-based version of our command-line tool CellNet. Learning and generalizing from this experience, we have devised a lightweight framework, Radiator, to facilitate deploying bioinformatics tools as web applications. To achieve reproducibility, usability, consistent accessibility, throughput, and cost-efficiency, Radiator is designed to be deployed on the cloud. Here, we describe the internals of Radiator and how to use it.

**Availability and Implementation:** Code for Radiator and the CellNet Web Application are freely available at https://github.com/pcahan1 under the MIT license. The CellNet WebApp, Radiator, and Radiator-derived applications can be launched through public Amazon Machine Images from the cloud provider Amazon Web Services (AWS) (https://aws.amazon.com/).

## Introduction

We recently published the RNA-Seq-compatible command-line version (Radley et al., 2017) of CellNet (Cahan et al, 2014), a computational platform for assessing the derivations of stem cell fate engineering protocols. To broaden accessibility to RNA-Seq CellNet, we developed the CellNet WebApp (https://github.com/pcahan1/CellNet_Cloud), a cloud-based web application which performs both raw data preprocessing and CellNet analysis and has additional functionality for analyzing human iPSC disease modeling studies (Fig. 1a, 1b).

**Figure 1.**
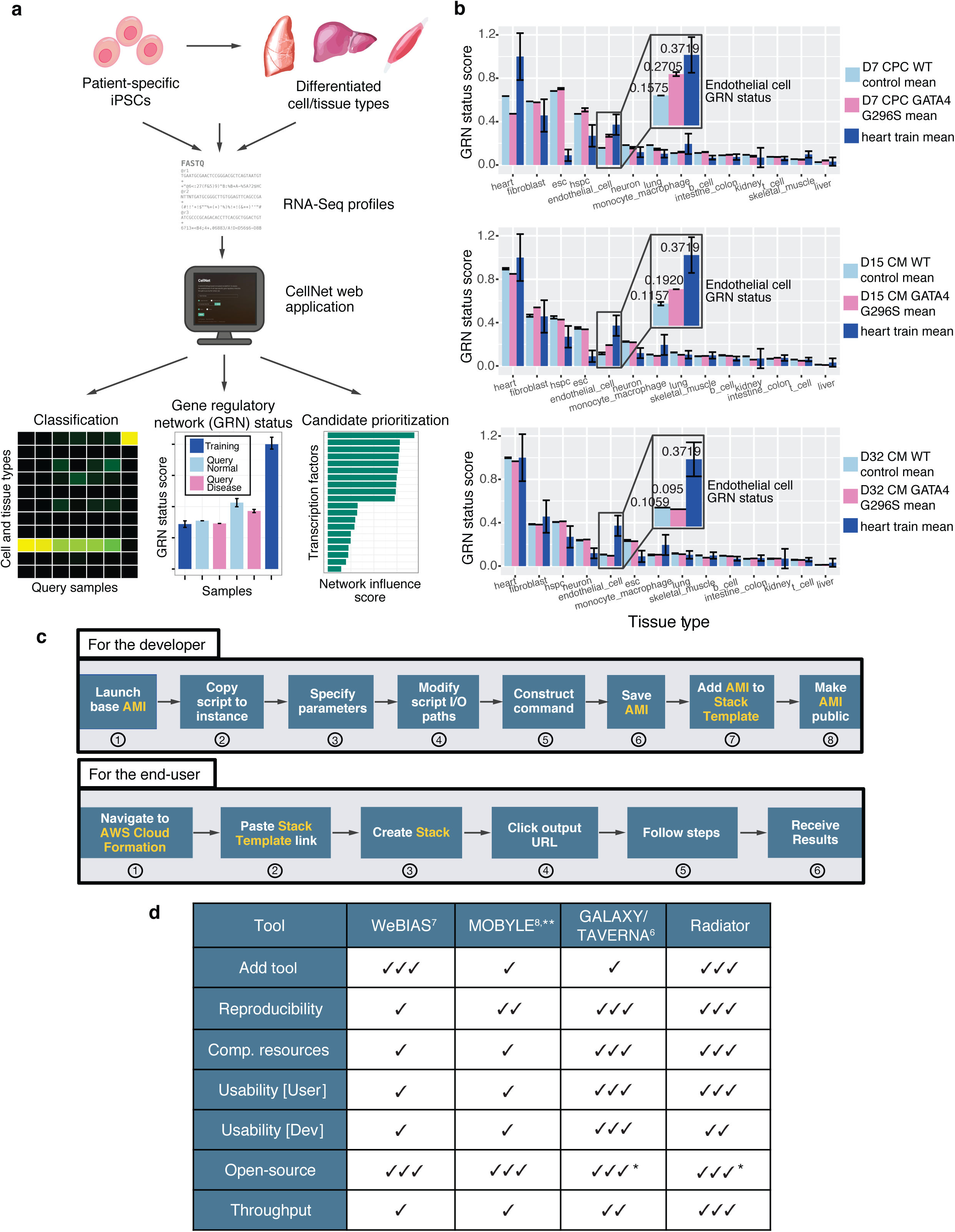
CellNet WebApp with iPSC disease modeling analysis capabilities, Radiator framework. **(a)** The CellNet WebApp performs both preprocessing to generate expression matrices and CellNet analysis to generate three readouts of cell identity: 1) a classification heatmap 2) the gene regulatory network (GRN) status, and 3) the Network Influence Score (Cahan et al, 2014; Radley et al., 2017) (Supplementary Information). **(b)** GRN statuses from an iPSC-disease modeling study of a cardiomyopathy caused by fate-altering mutation GATA4 G296, which results in upregulation of the endothelial GRN (Ang et al., 2016). GRN statuses for Day 7 (D7) iPSC-derived cardiac progenitors (top); D15 iPSC-derived cardiomyocytes (middle); and D32 iPSC-derived cardiomyocytes (bottom). Endothelial GRN status is higher and heart GRN status lower in the mutant than WT for D7 and D15 (Supplementary Fig. 1). **(c)** Procedure for developers to use Radiator framework to build web application; procedure for end-users to use web application. Orange text refers to AWS-specific terms (Supplementary Information). **(d)** Feature comparison for web platforms used in bioinformatics analysis. *Compute resources through cloud providers charge on a per-use basis. **Legacy project, no longer supported.

Developing this application highlighted several generic barriers to deploying compute-intensive bioinformatic pipelines. Users struggle with customizing tools for institution-specific compute resources and reconciling incompatible operating systems, software versions, input parameters, and genome index versions, which all contribute to issues with reproducibility. Developers also struggle with optimizing usability and consistent accessibility and supporting the cost of a compute-intensive tool (Beaulieu-Jones and Greene, 2017).

Thus, in developing the WebApp, we sought to ensure that the tool would (i) be cost effective for the developer, (ii) be elastic for the user, (iii) be usable, (iv) allow for resource-intensive computations, (v) encourage reproducible analysis, and (vi) be widely accessible (Langmead and Nellore, 2018). By deploying the application through a cloud provider, we were able to meet criteria (i-v). The burden of maintaining a server is obviated, as the cost of launching and using the application is shifted to the end-user on a per-use basis. Reproducible analysis is also furthered because all required software and libraries are fixed on each cloud ‘image’ on which the tool is deployed. Moreover, by developing a web-based interface we were able to meet the accessibility criterion (vi).

Recognizing that barriers to meeting these criteria would be encountered by any academic group seeking to deploy compute-intensive bioinformatic pipelines, we developed Radiator, a generalized, lightweight framework backed by the cloud-provider Amazon Web Services (https://github.com/pcahan1/Web_Framework) for bioinformaticians to easily deploy command-line tools as accessible web applications.

## Materials and Methods

Radiator is built with PHP, HTML5, CSS, JavaScript, JSON, and AWS Cloud Formation. Radiator utilizes a Model-View-Controller (MVC)-like architecture (Bucanek, 2009), an organizational paradigm that separates the logic of code into the categories of front-end appearance (views), back-end/server-side software (controllers), and data (models). CellNet is built with R, Salmon, cutadapt, and HISAT2, and the CellNet WebApp is built as with the Radiator framework.

CellNet and Radiator are hosted on publicly available Amazon Machine Images (AMIs) through AWS (Supplementary Information). We have configured the AMIs with requisite software and libraries preinstalled, which makes standardized and reproducible analysis feasible.

## Usage

The CellNet WebApp is activated on a per-use basis. Using the application requires only navigating to AWS Cloud Formation through the user’s AWS account, launching the specified CellNet AWS Stack, then following the simple and user-friendly interface to upload RNA-Seq data and study metadata. A step-by-step tutorial for using the CellNet WebApp is available (Supplementary Information).

Deploying an application through Radiator requires basic knowledge of Linux, virtual servers and machines, and web development. Within the Radiator framework, the developer modifies parameters and scripts to construct commands that will be elicited by the user through the web application. The end result is the configuration of an AWS AMI that interfaces with the user through a Cloud Formation web application, similarly to the CellNet WebApp. Developer and user workflows are detailed in Fig. 1c, and a tutorial for deploying a bioinformatics tool using Radiator is also available (Supplementary Information).

## Results and Conclusions

The CellNet WebApp has added functionality for analyzing iPSC disease modeling studies, especially with regards to the detection of cell-fate-altering mutations and disrupted transcriptional cascades in disease. In a study of a genetic cardiomyopathy (Ang et al., 2016a), CellNet was able to detect an aberrant endothelial signature in diseased iPSC-derived cardiomyocytes as compared to the signature seen in healthy controls (Fig. 1b, Supplementary Fig. 1).

We compare Radiator features to those of similar web platforms, including The Galaxy Project, WeBIAS, and Mobyle (Afgan et al., 2010; Daniluk et al., 2015; Goecks et al., 2010; Néron et al., 2009) (Fig. 1d, Supplementary Information).

The CellNet WebApp is a user-friendly tool for preprocessing and assessing RNA-Seq data from cell fate engineering experiments. We have generalized the WebApp framework as Radiator for bioinformaticians to distribute compute-intensive bioinformatics tools as web applications, while facilitating reproducible analysis, accessibility, and cost-efficiency for the developer and the end-user.

## Acknowledgements

We thank members of the Cahan Lab for helpful comments on the manuscript.

