## Supplementary Information for "Radiator: a cloud-based framework for deploying re-usable bioinformatics tools"

#### Supplementary Figures

**Supplementary Figure 1 I CellNet training and analysis of fate-altering mutations in disease.**

**(a)** Classifier performance heatmap of cross-validation samples from human RNA-Seq data. Each column represents an RNA-Seq sample; each row represents the binary random forest classifier for a cell/tissue type; heatmap color represents the score produced by the given classifier for the given sample. We trained CellNet using over 1000 publicly available primary non-diseased human RNA-Seq data samples across 103 studies and 14 cell/tissue types. The CellNet Web Application uses the CellNet processor object produced using this training data (Cahan et al, 2014; Radley et al., 2017).

**(b)** Precision-recall curves for each CellNet cell/tissue type-specific classifier. Each point represents the precision and recall at a different score threshold requirement for a sample to be classified as the given cell/tissue type.

**(c)** Classification heatmap for an iPSC disease modeling study of a genetic cardiomyopathy by Ang, et al. 2016 (GSE85623). This study generated wild type and patient-specific iPSC-derived cardiomyocytes to examine the fate-altering mutation GATA4 G296, which disrupts interaction with TBX5 and allows TBX5 to upregulate endothelial genes inappropriately.

**(d)** Embryonic stem cell (esc), endothelial cell, and heart GRN statuses for each treatment group in the study. To highlight the ability of CellNet to identify cell fate-altering mutations in disease, we asked whether CellNet would be able to detect the aberrant endothelial GRN. For D7 and D15 iPSC-derived cardiac progenitors and cardiomyocytes, the endothelial GRN status is indeed higher in the mutant than in the wild type. For these samples and the D7 ESC-derived GATA4 siRNA knockdown, the heart GRN status is also lower in the mutant than in the wild type. This disease modeling analysis functionality is available in the CellNet Web Application.

### Supplementary Information

#### Box 1: Glossary of Terms

**Amazon Web Services (AWS)*** **-** Amazon Web Services (AWS) is a suite of cloud-based resources that includes data storage (using the **Simple Storage Service**, or **S3**) and computation (using **Elastic Compute Cloud**, or **EC2**). AWS provides access to high-performance computing resources without the overhead of setting up and maintaining computer clusters locally. Sign up for AWS at https://aws.amazon.com/console/. Instructions for getting started with AWS can be found at http://docs.aws.amazon.com/AWSEC2/latest/UserGuide/get-set-up-for-amazon-ec2.html. AWS charges users based on increments of services used. For example, the c4.8xlarge
instance type currently costs $1.591 per hour.

**Access ID & Secret Access Key*** **-** Security credentials (provided during the AWS sign-up process) for performing account-specific tasks. E.g., must be input as parameters on the CellNet Web Application if user opts to access RNA-Seq FASTQ files from S3.

**Elastic Compute Cloud (EC2)*** - AWS's cloud computing service. Further documentation and tutorials available at https://aws.amazon.com/ec2/.

> **Amazon Machine Image (AMI)*** **-** A preserved virtual computer state with specific software and libraries preinstalled, which makes standardized and reproducible analysis feasible.

**Instance –** A running instantiation of a machine image, for which you can specify operating system and hardware requirements, e.g. RAM, CPU, storage. Changes made to the instance state while running the instance will not modify the AMI; only saving an AMI or creating a new AMI will preserve the state.

**Simple Storage Service (S3)*** - AWS's cloud storage service. Further documentation and tutorials available at https://aws.amazon.com/s3/. If user has more than 4GB of RNA-Seq data to be analyzed through the CellNet Web Application, storing them on S3 and providing the path to the Web Application is highly recommended.

**Cloud Formation*** - AWS's service for creating, organizing, and deploying a collection (**Stack**) of AWS resources to power an application. Further documentation available at https://aws.amazon.com/cloudformation/.

**Stack Template* -** A .JSON or .YAML document that defines the specific resources (e.g. AMI, database, instance type, web server security group) to be included in a Stack.

**Stack* -** A collection of AWS resources managed as a single unit. Essentially an instantiation of a Stack Template. The CellNet Web Application is deployed as a Stack.

**tar –** A Linux command for grouping or 'packaging' files together in an 'archive.'

**gzip –** A Linux command for file compression.

*Term specific to AWS

#### CellNet Outputs and Methods

CellNet analysis generates three readouts of cell identity: 1) a classification heatmap indicating how indistinguishable a query sample expression profile is from each of the reference cell/tissue types; 2) the gene regulatory network (GRN) status, a metric of the extent to which a cell/tissue-specific GRN is established in a query sample; and 3) the Network Influence Score (NIS), a list of transcription factors scored by the probability that their expression modulation would improve the desired fate change (Cahan et al, 2014; Radley et al., 2017).

For cell/tissue type classification, binary classifiers were trained for each individual cell/tissue type using the random forest algorithm. An overall gene regulatory network (GRN) was first reconstructed across all cell and tissue types. Cell/tissue type-specific GRNs were then identified by splitting the overall GRN into densely interconnected subnetworks through community detection, then attributed to a specific cell/tissue type through gene set enrichment analysis. More comprehensive methods for CellNet classifier training and validation, gene regulatory network reconstruction and validation, cell/tissue type-specific subnetwork identification, and computation of GRN status and Network Influence Score (NIS) have been described previously (Cahan et al, 2014; Radley et al., 2017).

The CellNet web application uses the following cnProc (Trained CellNet Processor) objects:

Mouse: cnProc_MM_RS_Oct_24_2016.rda
Human: cnProc_HS_RS_Jun_20_2017.rda

described in further detail at (https://github.com/pcahan1/CellNet).

### CellNet Web Application Walk-through Tutorial

The CellNet Web Application (https://github.com/pcahan1/CellNet_Cloud) delegates compute-intensive tasks to Amazon Web Services (AWS) compute resources. To fully utilize all features of the web application, users must have the following: an AWS account, username and password, AWS Access ID and Secret Access ID, and account permissions to access to EC2, S3, and Cloud Formation (Box 1).

Below is an outline of the steps needed to run the web application:

1. Login to AWS.
2. Select the Cloud Formation service from the main menu:

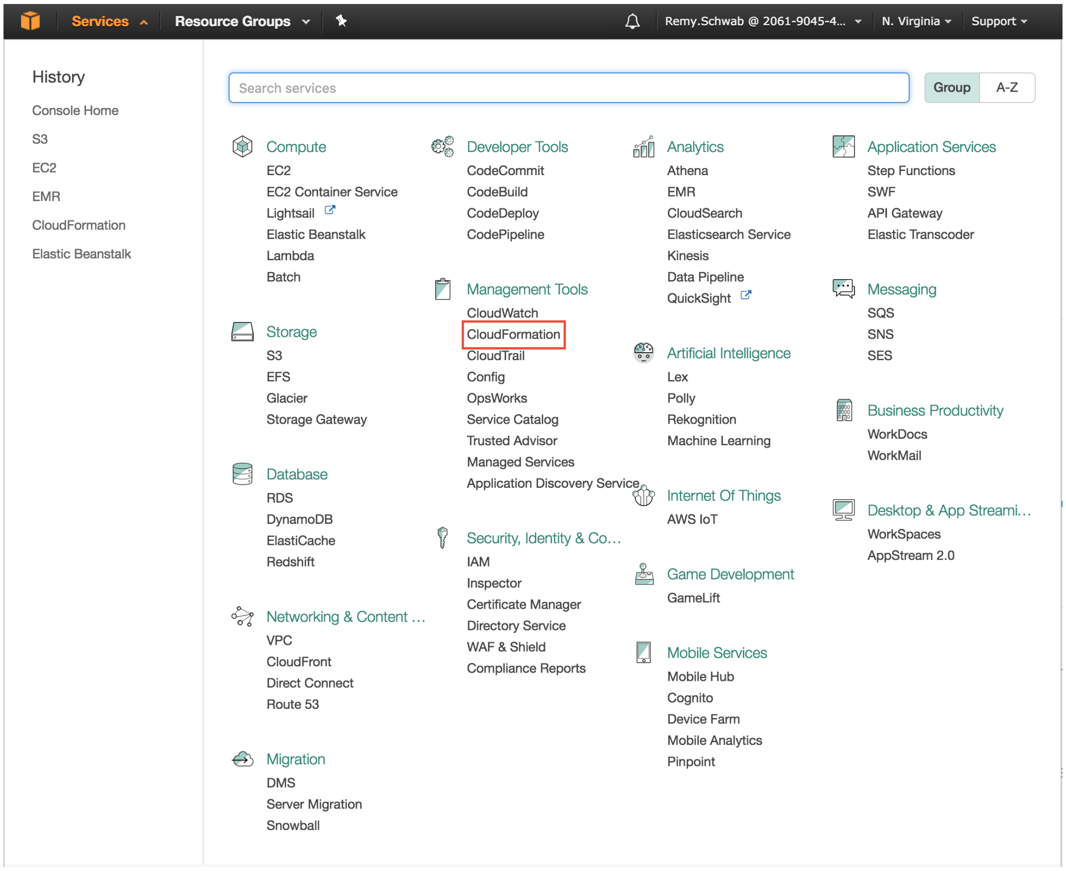

3. Click ‘Create New Stack’:

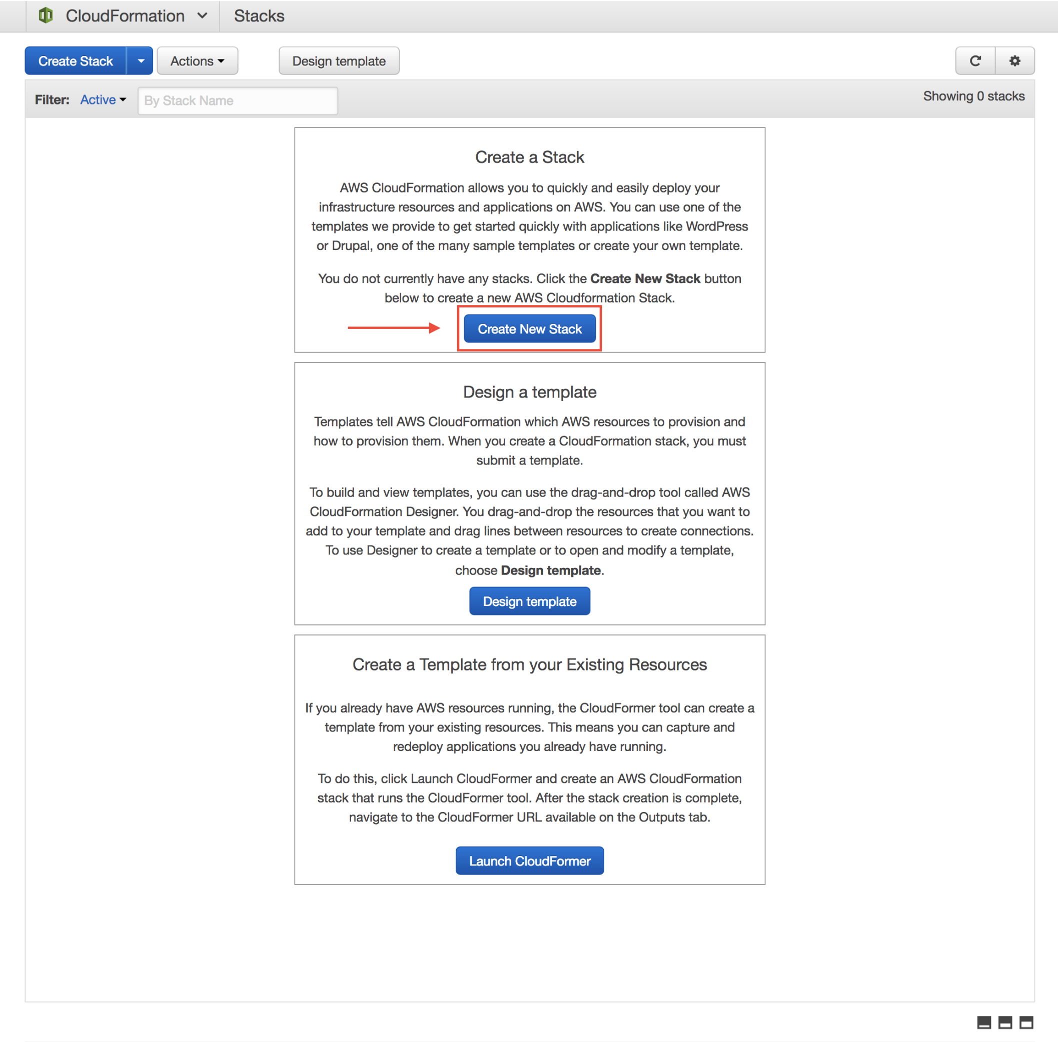

4. Select 'Specify an Amazon S3 template URL' and paste the link to the CellNet Stack Template: https://s3.amazonaws.com/cahanlab/remy.schwab/Stack_Templates/CellNet_publicStackTemplate.json

5. Name your Stack.

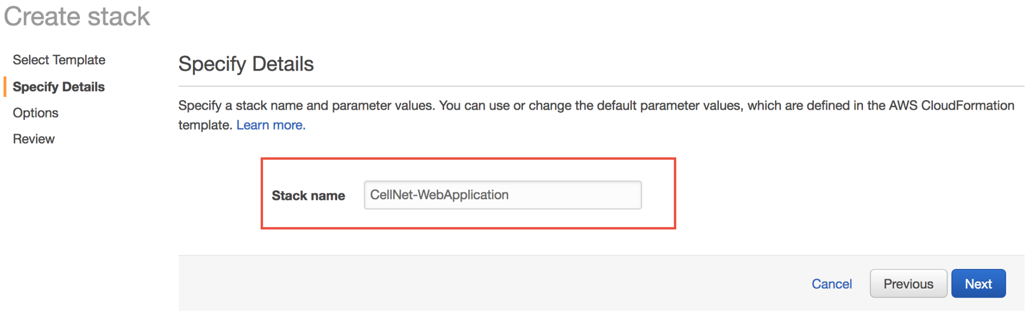

6. Skip the Options Page.
7. Review the details and hit ‘Create.’ This will launch an **EC2 instance** that is running the CellNet Web Application **AMI**.

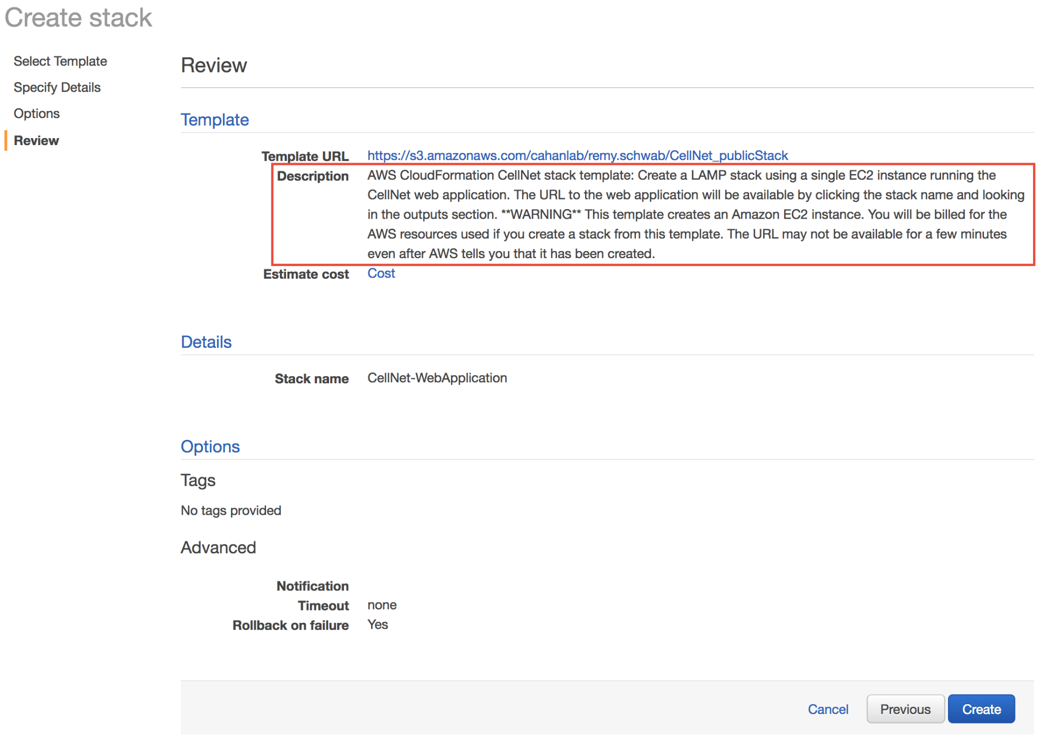

8. The instance should take about 5 minutes to initialize, but this can vary.

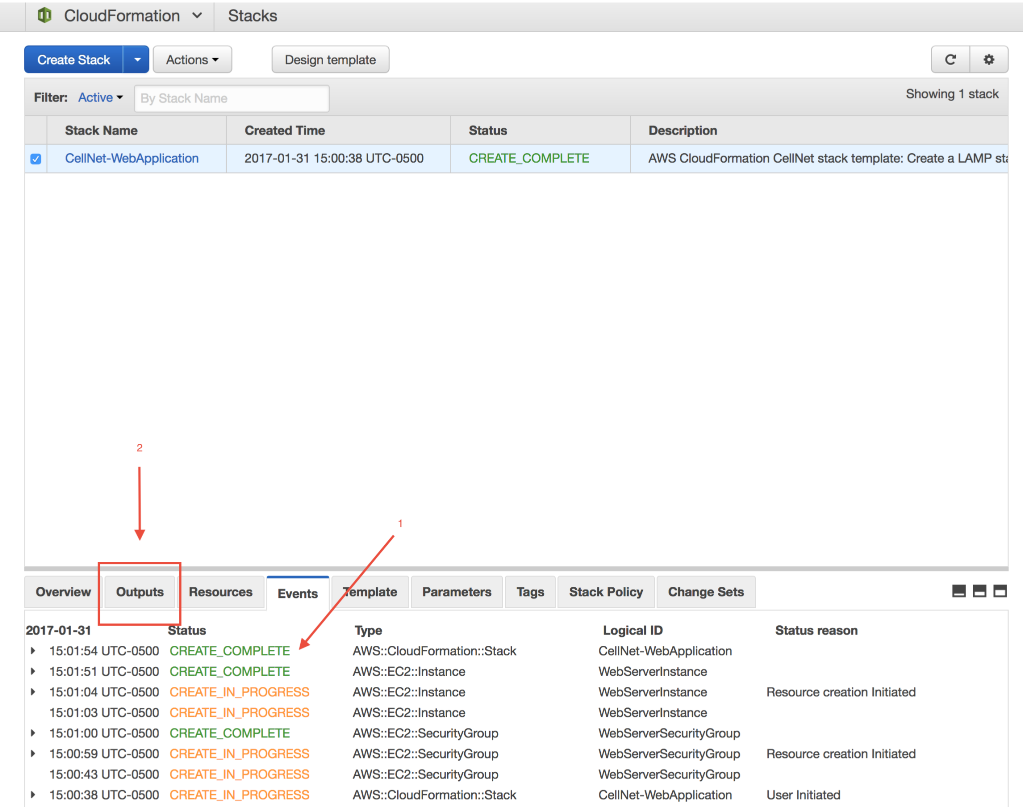

9. Once the instance is ready, a URL will be available under the ‘Outputs’ tab. Navigate to this URL in another window of your browser.

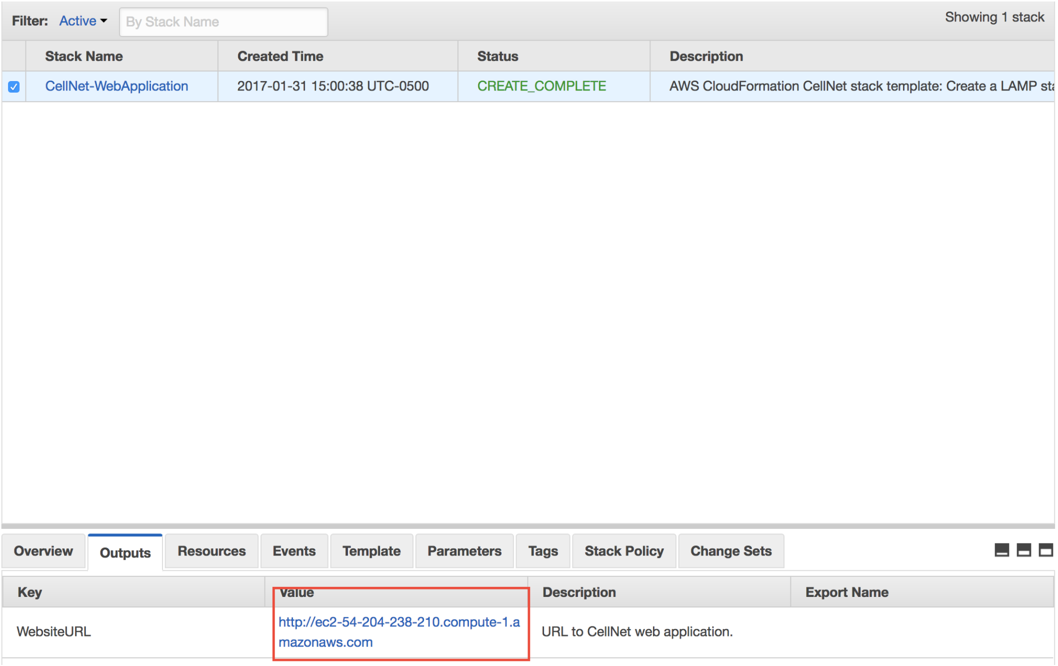

10. The Front Page
    a. Input your email address. Results will be emailed as an attachment.
    b. Upload sequencing files directly as a compressed archive OR specify where they are stored on AWS S3:

    The required format for uploading FASTQ files from a local machine is a **gzip**ped **tar** archive. This means that the uncompressed FASTQ files are compressed on the command line using a command like the following:

```
tar -cvfz new_archive_name.tgz fastq_folder_to_compress
```

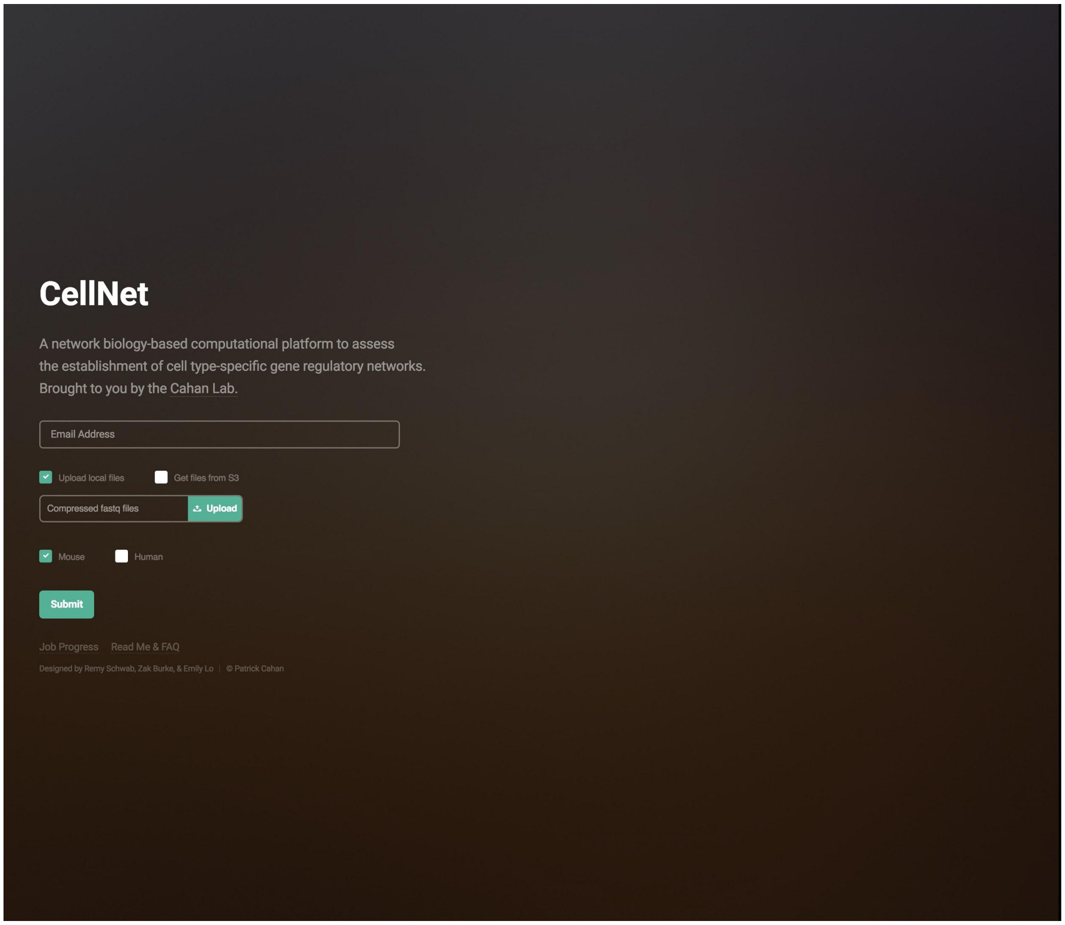

The local upload limit is capped at 4GB. If the resulting archive will be larger than 4GB (or if otherwise desired), files should first be stored on **AWS S3**, and a path to the files can be specified in the Web Application. These files must be stored in a folder containing only the FASTQ files.

Select ‘Get files from S3.’

Provide your AWS Access ID and Secret Access ID, and specify the S3 path to your files:

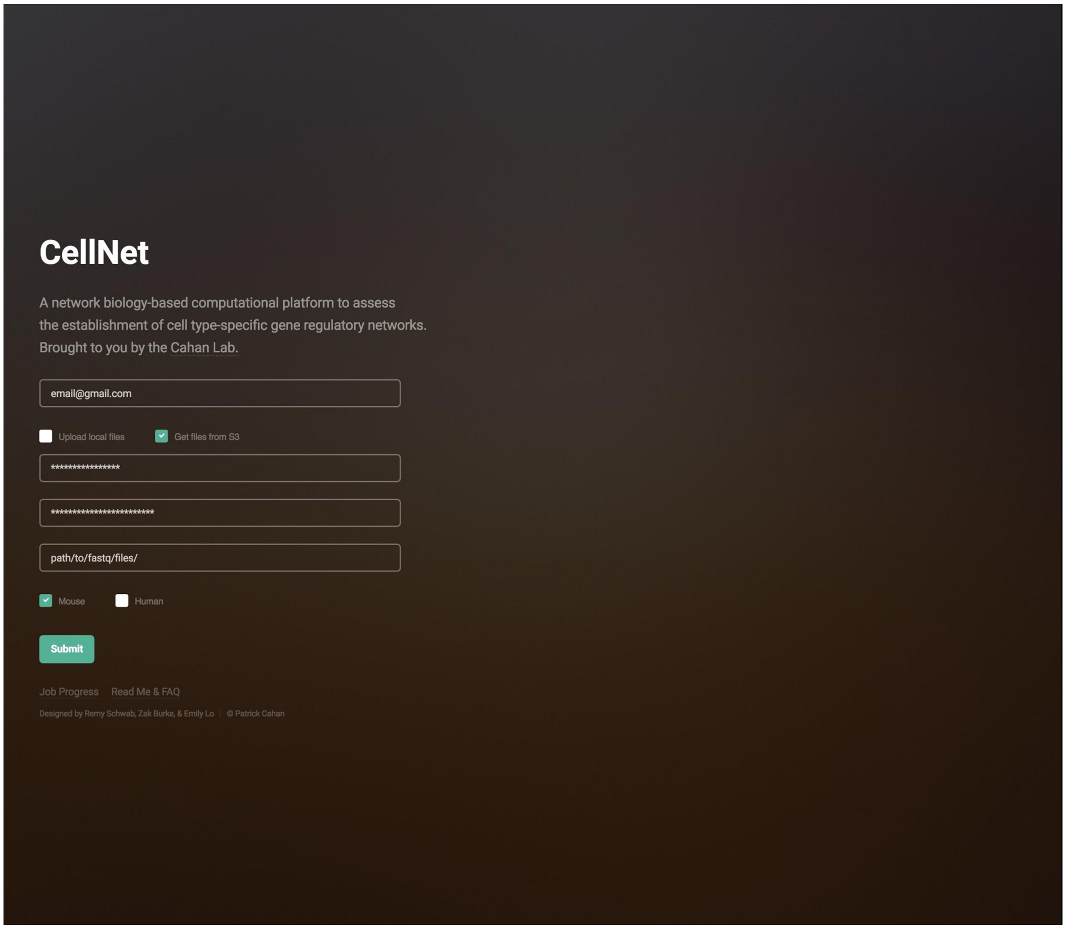

c. Specify species.
d. Submit. It may take several minutes to an hour to proceed to the next page depending on the size of the files, as both transfer and decompression is finished before the next step.

11.

a. Create the metadata table or upload your own (required metadata table format described in detail in Radley et al, 2016). Also specify starting and target cell types:

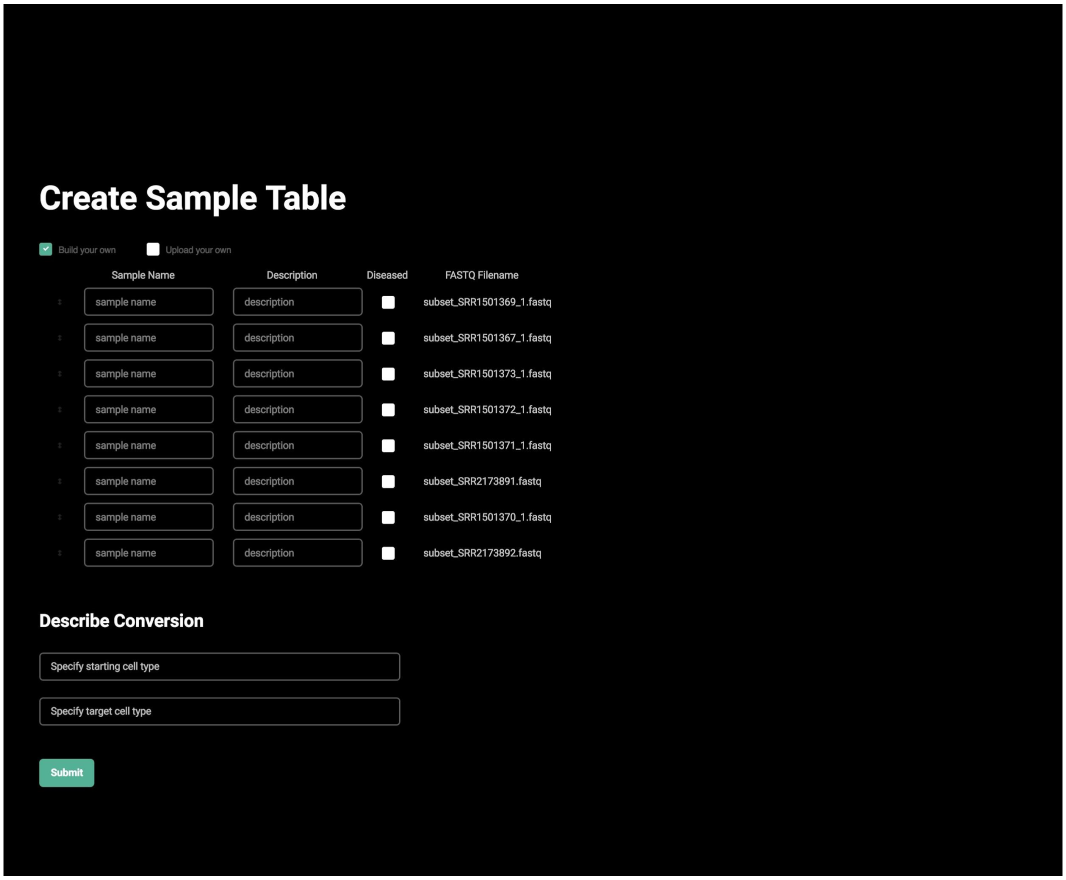
Create Sample Table
Build your own
Upload your own
Sample Name
Description
Diseased
FASTQ Filename
sample name
description
subset_SRR1501369_1.fastq
sample name
description
subset_SRR1501367_1.fastq
sample name
description
subset_SRR1501373_1.fastq
sample name
description
subset_SRR1501372_1.fastq
sample name
description
subset_SRR1501371_1.fastq
sample name
description
subset_SRR2173891.fastq
sample name
description
subset_SRR1501370_1.fastq
sample name
description
subset_SRR2173892.fastq
Describe Conversion
Specify starting cell type
Specify target cell type
Submit

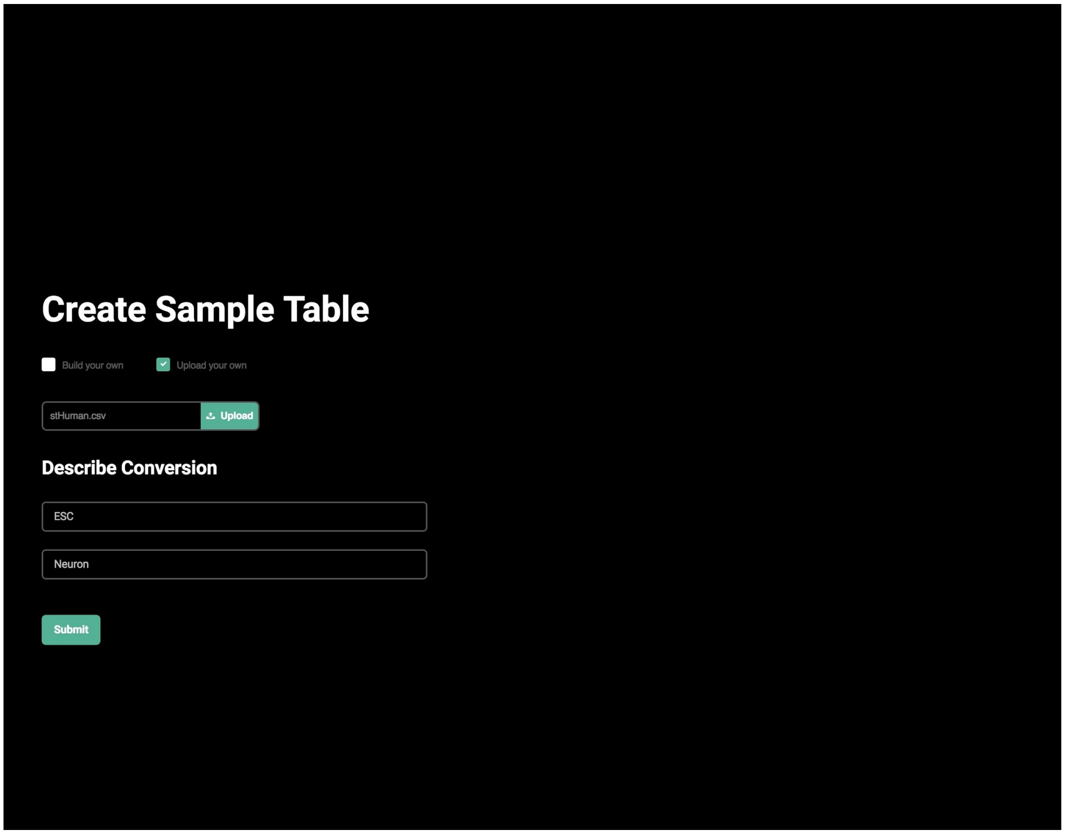
Create Sample Table
Build your own
Upload your own
stHuman.csv
Upload
Describe Conversion
ESC
Neuron
Submit

12. This screen displays the progress of your query submission.
    a. Several steps in this progress are slow, e.g. sequence read mapping and quantification. The entire process may take 30 minutes – several hours.

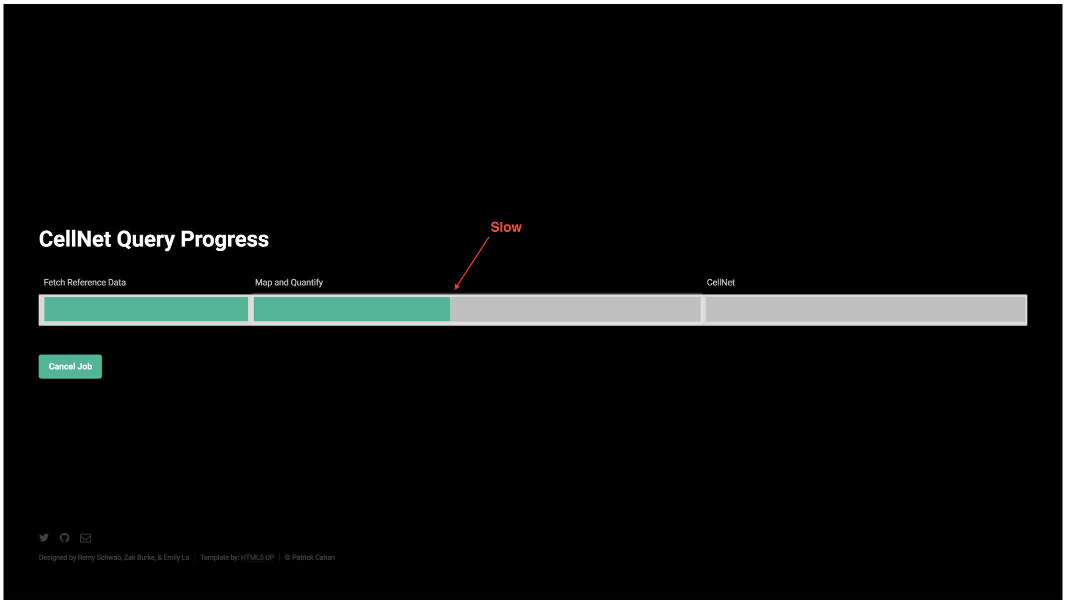

    b. This screen indicates analysis has finished:

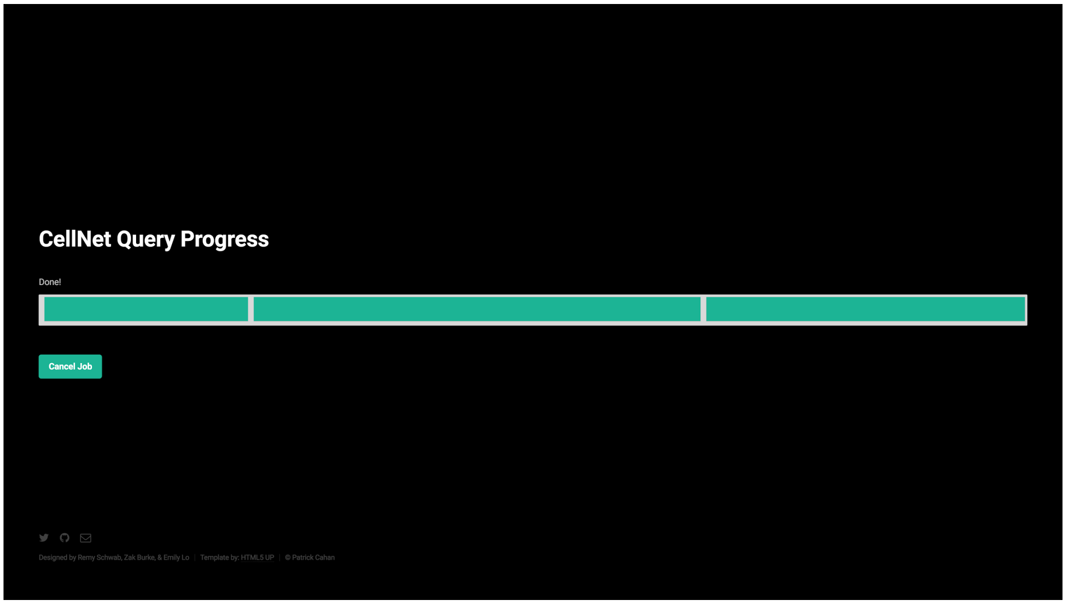

CellNet analysis is complete.

13. **IMPORTANT**: Once finished, make sure to delete the Stack (Actions > Delete Stack > Yes, Delete). This will terminate the running EC2 instance. AWS will continue to charge the user for computational resources used until the Stack is deleted.

#### Feature Comparison Criteria

The current leader in addressing these issues is the Galaxy Project, a widely used web-based platform that provides an accessible, transparent interface for running common bioinformatics tools reproducibly (Afgan et al., 2010; Goecks et al., 2010). While Galaxy is beginning to offer cloud computing functionality, it is geared more towards the user than the tool developer, making it difficult for an independent tool developer to make software available through Galaxy's cloud interface. Another previously published method, WeBIAS (Daniluk et al., 2015), focuses on increasing convenience of creating a front-end interface for command line tools by primarily abstracting away complexities of deploying a web application.

For each of the frameworks evaluated, below are descriptions of the criteria used in Figure 1d to assign feature scores. 3 – successfully addresses the criterion; 2 – attempts to address the criterion; 1 – may attempt to address the criterion, but unsuccessfully, or does not address the criterion.

**Add tool:** How feasible is it for an independent tool developer to make a new tool available?
**Reproducibility:** How conducive is the tool to reproducibility?
**Computational resources:** Does the tool support compute-intensive tasks?
**Usability [User]:** How user-friendly is the tool?
**Usability [Developer]:** How developer-friendly is the tool? Deploying a web application through Radiator requires basic knowledge of Linux, virtual cloud machines and servers, and web development concepts; therefore, we gave it a score of 2.
**Open-source:** Is source code for the tool publicly available? Is there an upfront cost to using the tool? Exception: compute resources through cloud providers have a cost, e.g. Galaxy's CloudMan Interface (Afgan et al., 2010) and the CellNet Web Application, which are both deployed through AWS.
**Throughput:** Does the tool facilitate high-throughput analysis through a cloud provider? Though the Galaxy CloudMan Interface (Afgan et al., 2010) does provide access to high performance computational resources, it does not allow persistent access to an independent tool through the cloud.

### Radiator Walk-through Tutorial (For the Developer)

Radiator (https://github.com/pcahan1/Web_Framework) is a web framework that allows you to deploy your bioinformatics command line tool through a cloud provider, facilitating flexibility, computationally efficiency, cost-effectiveness, and above all, reproducibility.

The following walkthrough assumes basic knowledge of Linux, virtual servers and machines, and web development concepts. **It will demonstrate how to adapt our base framework for a simple example application.** To help familiarize with AWS services, we have also provided further information about AWS virtual machines and other resources (Box 1).

We have written a Python 'down-sampling' script as an example of a bioinformatics tool one might want to deploy. The script takes a random sample of reads from a FASTQ file; this is useful when fewer reads than may be present in a FASTQ file are sufficient for a downstream analysis tool.

#### Architecture overview:

Radiator utilizes a Model-View-Controller (MVC)-like architecture. This is an organizational paradigm that separates the logic of code into the categories of front-end/user-side appearance (views), back-end/server-side software (controllers), and data (models).

In the Radiator framework, directories are organized under the MVC paradigm as such: HTML files are found under 'views', server-side scripts under 'controllers', and data under 'models'.  Supplemental CSS and Javascript files specifying additional front-end behaviors are found under ‘assets.’ These directories are stored in the server's filesystem under **/var/www/html** (See Radiator Directory Structure below).

#### Radiator Directory Structure:

AMI Name: Cahan-Lab Application Framework Base Image
AMI ID: ami-5dd1db27

```
/var/www/html/
|
├── assets
│   ├── css
│   │   ├── index.php
│   │   ├── main.css
│   │   └── upload-file.css
│   └── index.php
├── controllers
│   ├── Algorithm
│   │   └── index.php
│   ├── index.php
│   └── run.php
├── index.php
├── models
│   ├── input
│   │   ├── subset_SRR1501367_1.fastq.gz
│   │   ├── subset_SRR1501368_1.fastq.gz
│   │   ├── subset_SRR1501369_1.fastq.gz
│   │   ├── subset_SRR1501370_1.fastq.gz
│   │   ├── subset_SRR1501371_1.fastq.gz
│   │   ├── subset_SRR1501372_1.fastq.gz
│   │   ├── subset_SRR1501373_1.fastq.gz
│   │   ├── subset_SRR2173891.fastq.gz
│   │   └── subset_SRR2173892.fastq.gz
│   └── output
│       └── compressed_output.tgz
└── views
    ├── front_page.php
    ├── index.php
    └── results_page.php
```

#### Launching and Adapting the Base Radiator Framework:

The base virtual machine is hosted by Amazon Web Services as an EC2 Amazon Machine Image (AMI). Currently it is only available in the US-East-1 region.

1. Select the Radiator AMI: Cahan-Lab Application Framework Base Image <AMI ID: ami-5dd1db27>
2. Specify desired instance type.
3. Specify security group. Make sure it allows for incoming traffic through ports 20 (SSH for shell access) and port 80 (HTTP for web access)
4. Launch instance.

#### The Front Page:

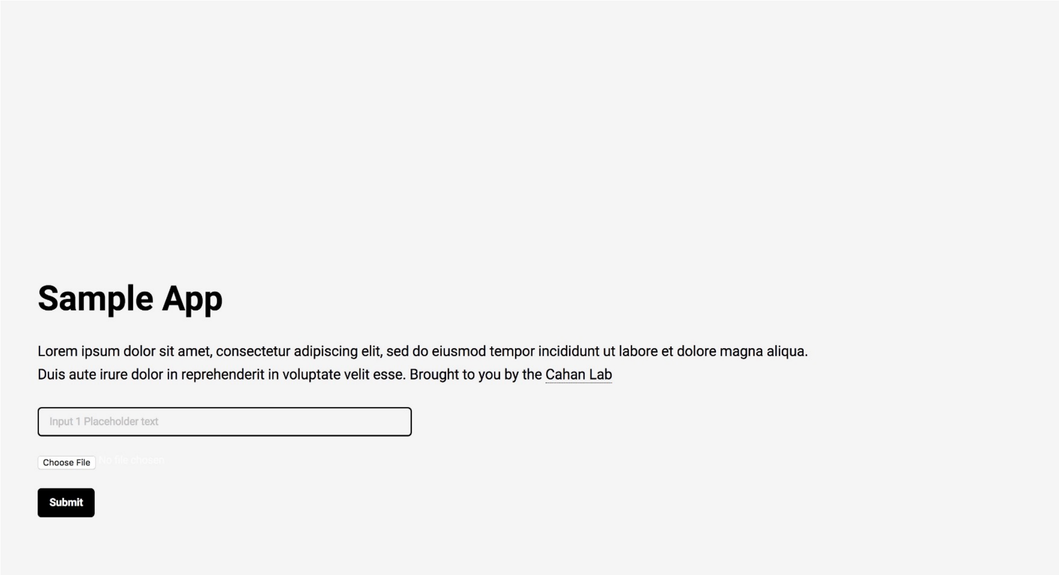

5. Modify the input parameters of the front page by editing front_page.php in **/var/www/html/views/**.

    Name your application and provide a brief description here.

    Add HTML form elements for specifying the user parameters necessary for analysis.

    Modify application appearance in **/var/www/html/assets/css/**main.css and upload-file.css.

6. For the purposes of the down-sampling application, we will add an html form element to specify the number of reads desired.
7. Add the following HTML tag to the front page:

```
<input type="text" name="parameter1" placeholder="Input Read Depth [Default 5000000]"/>
```

8. Now your front page should look similar to the following:

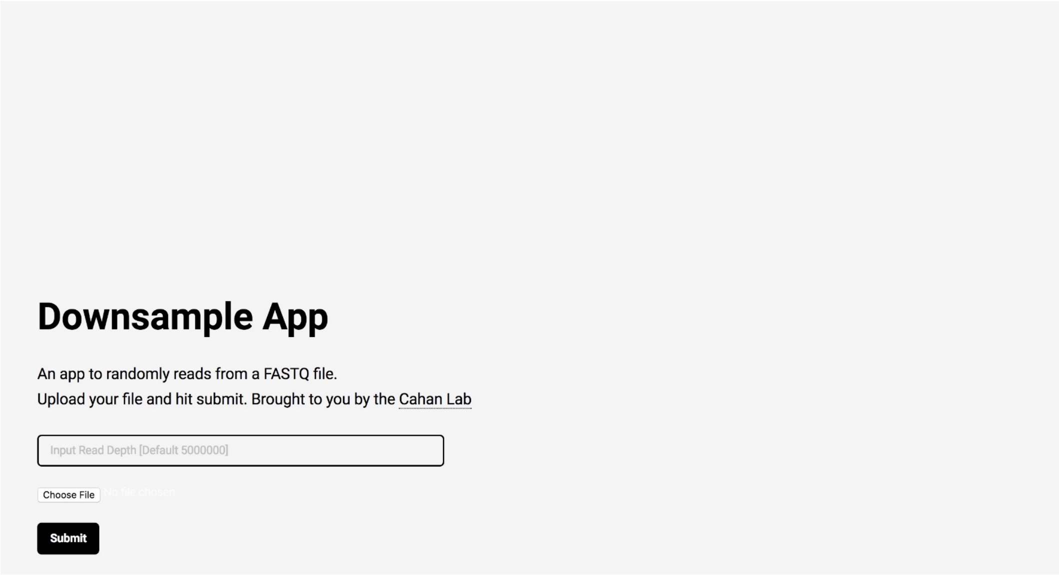

#### Controllers:

9. Add your executable (in this case, the down-sampling executable) to its proper location using scp. Your command should look something like this:

   scp -i keyname.pem downsample_file:/var/www/html/controllers/Algorithm

10. Assign the proper permissions so that Apache can execute it:

    sudo chown apache downsample_file
    sudo chmod +x downsample

The run.php script collects parameters specified by the front-page form so that you can build a Linux shell command that will feed input into your script.

To build the command, change the block

```
if(isset($_POST['parameter1'])) {
    $parameter1 = (int)$_POST['parameter1'];
    #$command = "/var/www/html/controllers/Algorithm/EXECUTABLE_NAME";
}
else {
    #$command = "/var/www/html/controllers/Algorithm/EXECUTABLE_NAME";
}
```

to

```
if (move_uploaded_file($_FILES['data']['tmp_name'], $target_file)) {
## CHECK IF A PARAMETER IS SPECIFIED
    if(isset($_POST['parameter1'])) {
        $parameter1 = (int)$_POST['parameter1'];
        $command = "/var/www/html/controllers/Algorithm/downfile -n $parameter1 $target_file";
    }
    else {
        $command = "/var/www/html/controllers/Algorithm/downfile $target_file";
    }
    exec($command);
}
```

#### Setting I/O Paths:

Ensure that the executable is writing its output to the correct location. In the down-sampling script, replace this line (LINE 40):

```
with open("subset_"+fname, "w") as output:
```

with

```
with open("/var/www/html/models/output/subset_"+fname, "w") as output:
```

#### Make Application Available through Cloud Formation:

11. Save new AMI based on current instance state.
12. Make the AMI public.
13. Map your Stack Template to launch an instance of your AMI.
14. Make the link to your Stack Template publically available.

#### Launching the Application (End-user):

This process is similar to launching the CellNet web application, detailed in the **CellNet web application walk-through tutorial** above.

15. Navigate to Cloud Formation.
16. Create Stack by pasting link to Stack Template.
17. Use the output URL to open the web application and use as desired.
18. Finish and delete stack.
