## Supplementary figures and images for "Radiator: a cloud-based framework for deploying re-usable bioinformatics tools"

### Supplementary Figure 1

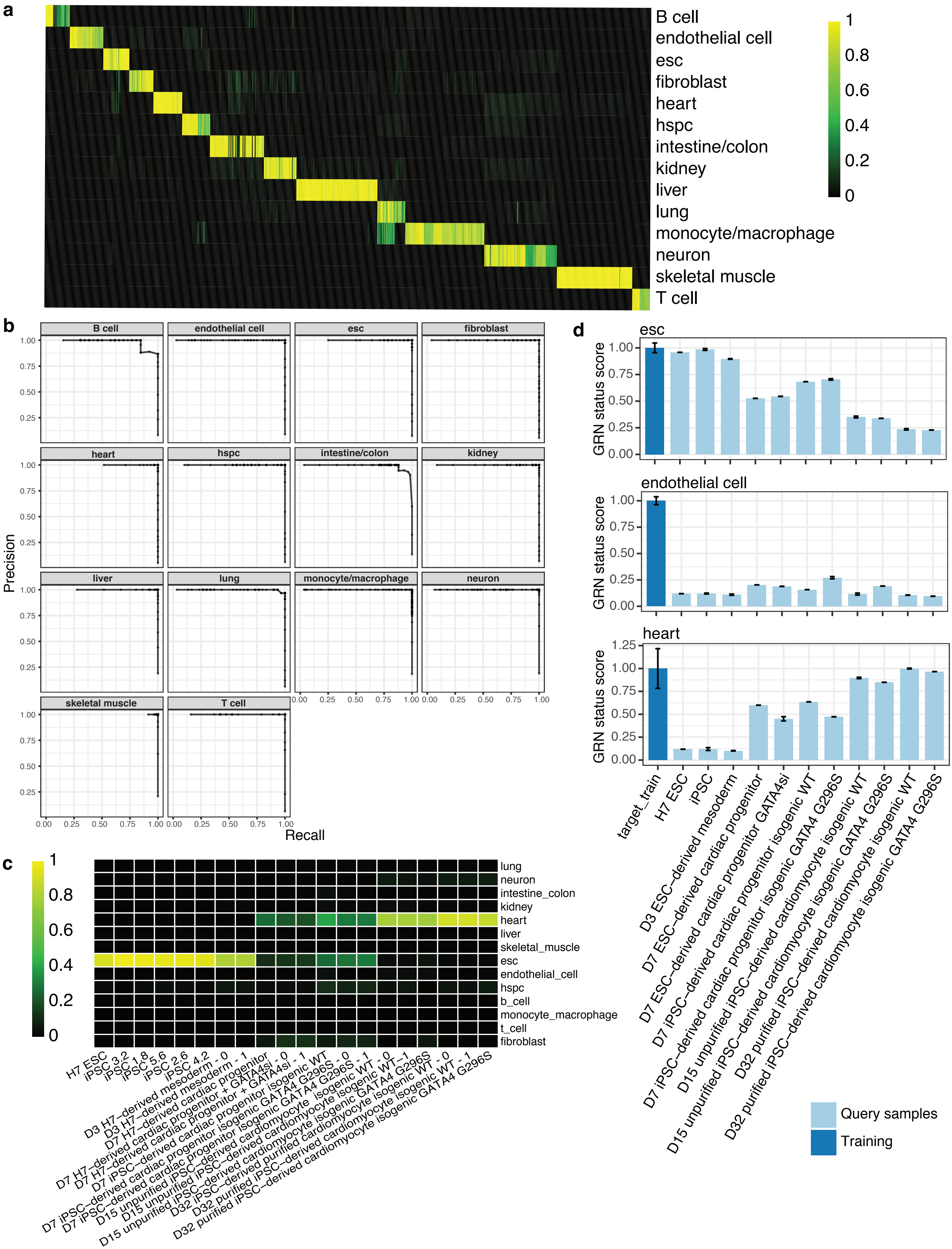
